# Non-epigenetic mechanisms enable short memories of the environment for cell cycle commitment

**DOI:** 10.1101/2020.08.14.250704

**Authors:** Yimiao Qu, Jun Jiang, Xiang Liu, Xiaojing Yang, Chao Tang

**Affiliations:** Center for Quantitative Biology and Peking-Tsinghua Center for Life Sciences, Academy for Advanced Interdisciplinary Studies, Peking University, Beijing 100871, China; School of Physics, Peking University, Beijing 100871, China

## Abstract

Cells continuously survey their environment in order to make fundamental decisions, including whether to divide, migrate, or differentiate. However, a fascinating phenomenon in biology is that cells often possess memory—they temporally integrate both past and present signals to make a reliable decision. Cellular memory manifests across different biological systems over different timescales, and a variety of underlying molecular mechanisms have been proposed. Here we investigate a non-epigenetic molecular mechanism underpinning how a single yeast cell can remember its recent environmental history to decide whether to enter the cell cycle. This “memories” is encoded by the phosphorylation level of the cell cycle inhibitor Whi5. G1 cyclin Cln3 senses environmental nutrient levels and promotes cell-cycle entry by phosphorylating and thus inactivating Whi5. We developed an optogenetic system whereby the nuclear localization of Cln3 can be rapidly and reversibly controlled by light. By monitoring cellular response to different temporal profiles of Cln3, we found that cell cycle entry requires the time duration of nuclear Cln3, supporting the model of “cellular memories”. Moreover, instead of the memory could last for the entire G1 phase as previously observed in glucose, we found Whi5 re-activates rapidly, with a similar half-time ∼ 12 min, in a variety of nutrient and stress conditions. Our results suggest yeast cell can shortly remember its recent environmental cues to decide whether to enter the cell cycle.

## Introduction

Many types of cells display memory: their present behavior is shaped by their exposure to past signals (Burrill and Silver, 2010; Csermely et al., 2020; Schaefer and Nadeau, 2015). These cellular memories manifest over different timescales in different biological venues, and multiple underlying molecular mechanisms have been proposed. Epigenetic memories (e.g., those encoded by DNA methylation or histone modifications) are among the most long-lived, as they can be stably inherited across cell divisions or even generations (Gaydos et al., 2014; Perez and Lehner, 2019). Other molecular means have been proposed for cells to record memories of varying lengths, including long-lived proteins (Doncic et al., 2011; Gao et al., 2018; Qu et al., 2019) or the ability of proteins to adopt extremely stable conformational states or form long-lived aggregates (Harvey et al., 2018), thus enabling the transmission of information across cell divisions. At the circuitry level, the hysteresis or memory effect of bistable systems has long been documented (Gardner et al., 2000; Monod and Jacob, 1961; Novick and Weiner, 1957), where the cell tends to persist in the past state despite environmental changes. On the other hand, the possibility of protein modification serving as short-term memory was less known, with the exception in *E. coli* chemotaxis in which the methylation level of the chemoreceptor reflects the recent concentration of chemo-attractant or repellant in the environment (Berg, 2000; Ma et al., 2009; Tu, 2013). Here we investigate a molecular mechanism underlying an emerging example of short-term cellular memory encoded by protein phosphorylation, namely how yeast record their recent environmental history in order to decide whether to divide in the present.

Yeast cells decide whether or not to enter the cell cycle by assessing multiple environmental signals, including the presence of nutrients, stressors, hormones and growth factors (Johnson and Skotheim, 2013; Jorgensen and Tyers, 2004). Cln3 (the homolog of Cyclin D in mammals), is a key protein used by yeast cells to assess whether nutrients in the environment are sufficiently rich in order to divide. Cln3 is a positive driver of the cell-cycle and can be construed as a nutrient sensor: it is transcriptionally and translationally upregulated in rich medium (Gallego et al., 1997; Hall et al., 1998; Parviz et al., 1998; Polymenis and Schmidt, 1997; Verges et al., 2007; Yahya et al., 2014). Moreover, Cln3 is a rapid nutrient sensor, as it is quickly upregulated by nutrients and also swiftly downregulated in their absence—both within minutes—enabling a fast response to changing external nutritional states (Cai and Futcher, 2013; Cross and Blake, 1993; Tyers et al., 1992; Wang et al., 2009; Yaglom et al., 1995). Cln3 forms a complex with Cdk1 (cyclin-dependent kinase 1) and promotes entry into the cell cycle: it phosphorylates and thus inhibits the cell-cycle inhibitor Whi5, leading to its nuclear export (de Bruin et al., 2004). Cells irreversibly commit to the cell cycle when ∼50% of Whi5 exits the nucleus (Doncic et al., 2011; Schmoller et al., 2015).

It has long been proposed that yeast cells continuously measure the concentration of Cln3, and when it reaches a certain threshold at any given time, they instantaneously commit to the cell cycle – the “instantaneous measurement” model whereby cells make snap decisions based on the current Cln3 level, i.e., “memory-less” (Jorgensen and Tyers, 2004; Schneider et al., 2004). However, our previous study found a negative correlation between Cln3 level and G1 duration, indicating that Cln3 signal in the nucleus is being integrated over time and the cell commits to cell cycle (Start) when Cln3-integration reaches a threshold value. The memory length was found to be comparable to the entire G1 length in SD medium, i.e., “entire-G1 memory” (Liu et al., 2015).

Here, by developing an optogenetic system where nuclear Cln3 activity can be fine-tuned by light rapidly and reversibly at the single-cell level, we showed that cell-cycle entry requires the time duration of nuclear Cln3 signal, consistent with the “memory” model. Furthermore, we found that, rather than integrating the environment cues over the entire G1, cells have a comparable short memory, about 12min, independent of the environmental conditions.

## Results

### Construction of the Cln3-controlled system using PhyB-PIF optogenetic module

Here, we sought to directly distinguish between two models for cell-cycle entry: the “instantaneous measurement” model vs. the “memory” model. One direct way is to check whether the cell cycle entry can be triggered by a transient Cln3 pulse in the nucleus. To generate this short pulse, we employed the PhyB-PIF optogenetic organelle-targeting system (Yang et al., 2013) to drive Cln3 in and out of the nucleus rapidly. PhyB associates with PIF in red light and they dissociate in infrared light, and both the association and dissociation are rapid, in seconds. Here, by using CAAX-tagged PhyB, we can tether the PIF fused Cln3 to the plasma membrane, thus out of the nucleus, in red light (the OFF state; Figure 1B). When exposed to infrared light, PhyB and PIF dissociate, and Cln3 can enter the nucleus (the ON state; Figure 1B). As fluorescence signals of wild-type Cln3 tagged with fluorescence proteins cannot be detected due to the fast and constitutive degradation of Cln3, here we used the stabilized mutant Cln3 (referred to as Cln3*) (Liu et al., 2015), which is an R108A substitution. This Cln3 mutation has been shown to not affect the capability to activate Cdk1 (Miller et al., 2005). To exclude the effects of other cell cycle inducers, we constructed this system in a *bck2Δ* background, where the role of Cln3 could be exclusively tested (Epstein and Cross, 1994).

**Figure 1.**
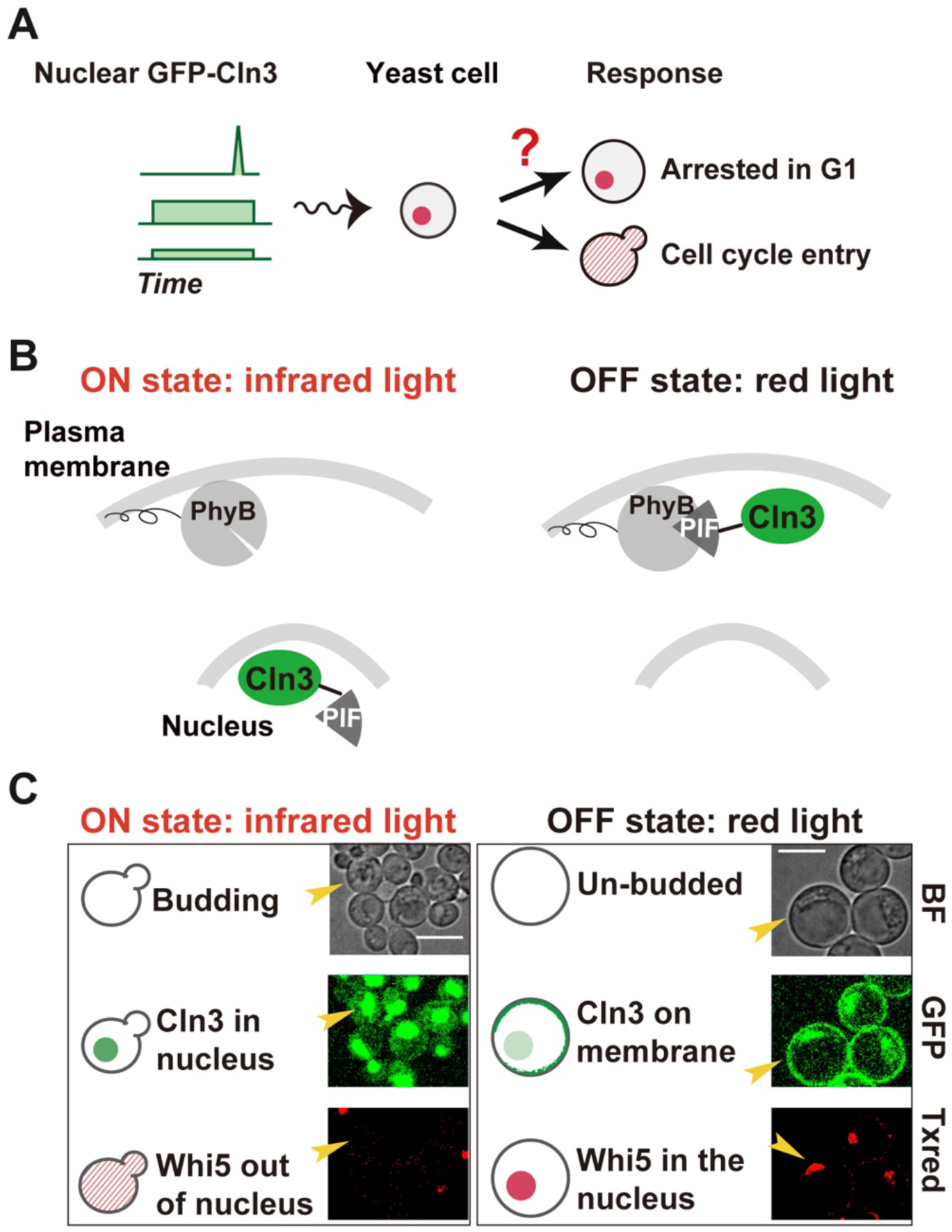
Control the nuclear Cln3 intensity in single cells using PhyB-PIF optogenetic module. (A) Schematic of hypothesis that cells process nuclear Cln3 dynamics to control cell cycle entry. (B) Schematic of constructing the nuclear Cln3-controlled system using the Phy-PIF optogenetic module. Cln3* is anchored to the plasma membrane when cells are exposed to the red light (OFF state). Cln3* stays in the nucleus when cells are exposed to the infrared light (ON state). (C) Different cell responses when exposed to infrared light (left panel) and red light (right panel). Yellow arrows indicated represented cells.

Then we demonstrated that this optogenetic system could be used to rapidly and reversibly control the nuclear localization of Cln3*. Firstly, when exposed to infrared light (ON state), yeast cells proceeded through the cell cycle normally compared to wild-type strain, while cells were arrested in G1 when kept in red light (OFF state), demonstrating that enough Cln3* control can be achieved using this system (Figure 1C and S1A). Arrested cells showed typical G1 state features: they did not bud and cell cycle inhibitor Whi5 remained in the nucleus (Doncic et al., 2011; Liu et al., 2015; Papagiannakis et al., 2017). Secondly, we tested the titratability of the nuclear Cln3*. We found the Cln3* level in the nucleus can be fine-tuned by changing the ratio of 650:750nm light intensity (Figure S1B). Thirdly, we measured how quickly the localization of Cln3* could be changed by switching light conditions. Both nucleus-to-cytoplasm and cytoplasm-to-nucleus translocation of Cln3* occurred rapidly (half-time ∼5 min) upon exposure to red and infrared light, respectively (Figure S1C). The longer timescale comparing to the association rate of PhyB and PIF may be due to the relatively slower nucleus-cytoplasm shuttling speed of Cln3.

### Cell cycle entry depends on the time duration of nuclear Cln3

To create a short pulse of nuclear Cln3, first yeast cells were arrested in G1 by red light, and subsequently they were exposed to a 5-minute infrared light pulse (Figure 2A). This sharp nuclear Cln3 pulse was unable to trigger Start (Figure 2C), even though its maximum nuclear Cln3 intensity (Cln3_max_) was significantly higher than the Cln3 intensity that were sufficient to trigger Start in cells constantly exposed to infrared light (ON state) (Figure 2B). Therefore, this result suggests that very high levels of Cln3 over a short timeframe cannot trigger Start, thus excluding the “instant measurement” model; rather it suggested that a prolonged increase of Cln3 is essential to trigger Start (i.e., the “time integral” model).

**Figure 2.**
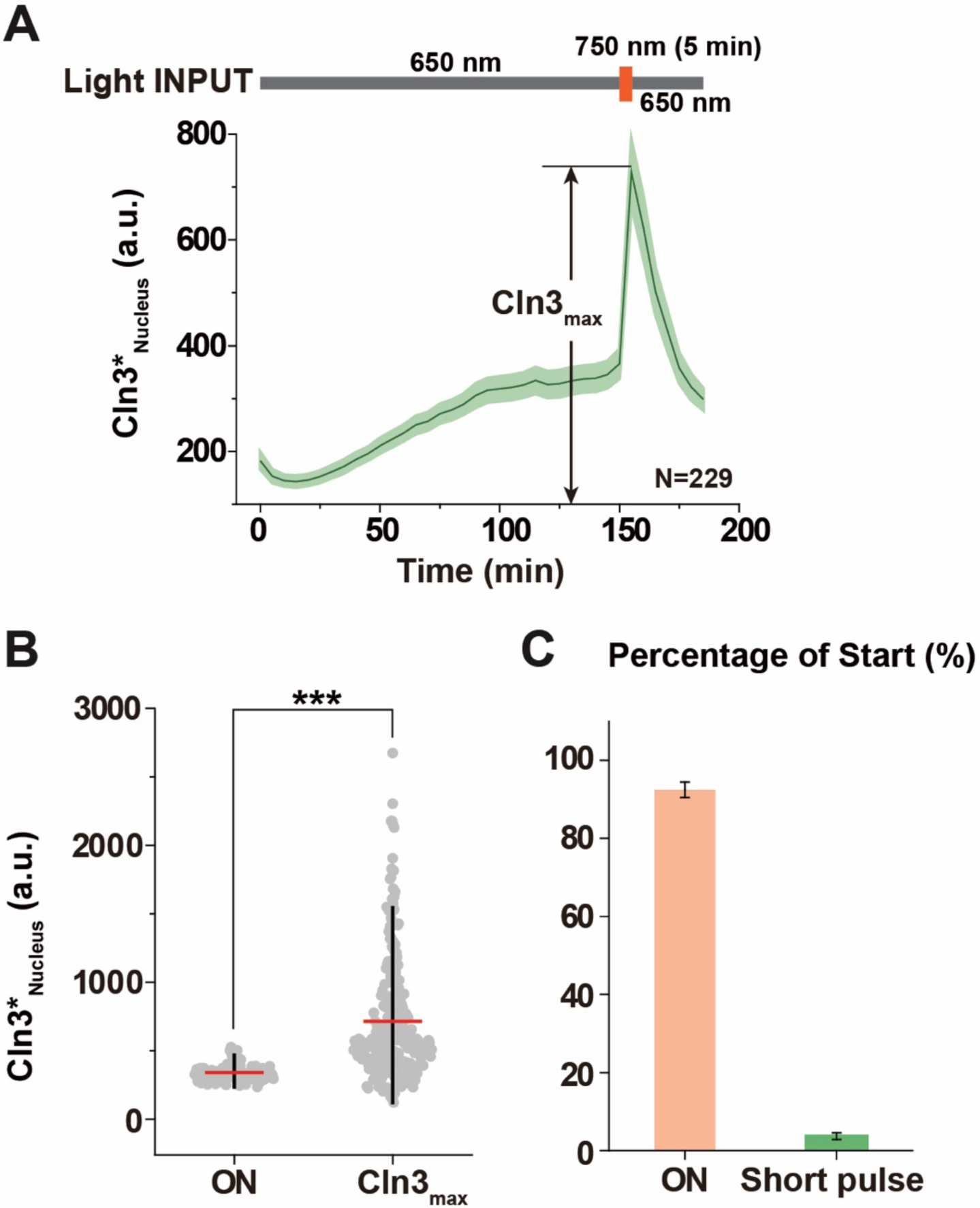
Cell cycle entry depends on the time duration of the nuclear Cln3 rather than its instantaneous intensity. (A) Real-time observation of the nuclear Cln3* intensity in cells (bottom panel) that are exposed to the indicated light input (top panel). Green curve, the mean nuclear Cln3* intensity. Shading, 95% confidence interval of the mean. See Methods for light power. (B) The maximum nuclear Cln3* intensity of the short pulse (shown in A) is higher than the averaged nuclear Cln3* intensity in G1 phase of cells exposed to the infrared light (ON state). Red bars, mean. Black bars, 25%-75%. ***, p value < 0.05. (C) The percentage of cells that passed through the Start (percentage of Start) point in response to the indicated light input shown in A.

We confirmed that the time duration of the Cln3 pulse is essential for triggering Start by using two different temporal strategies to increase nuclear Cln3 in cells (Figure S2). In the first strategy, cells were exposed to a sharp Cln3 pulse (which failed to trigger cell cycle entry), 30 min later followed by a longer Cln3 pulse with a lower intensity (Figure S2A). Most cells passed through Start only in response to the second, longer Cln3 pulse despite its lower intensity instead of the first high-intensity but short pulse. In the second strategy, to exclude the effects of the pulse sequence on triggering Start, we reversed the sequence of the two kinds of Cln3 pulses. Although the percentage of Start during the time of the sharp pulse increased from 1.1% to 15.5% after reversing (Figure S2A and S2B), the percentage of Start in response to the longer pulse with a lower intensity (67.5%) was still significantly higher than that in response to the sharp pulse. In summary, we showed that the temporal dynamics of Cln3 are critical to trigger Start, suggesting that cell cycle entry depends on the accumulation of Cln3 over time during G1, consistent with the “temporal memory” model (Liu et al., 2015).

### Whi5 rapidly re-enters the nucleus without CDK activity under various conditions

Next we studied how long the effects of Cln3 can last – it was previously proposed to span the whole G1 phase (Liu et al., 2015). Generally, Start is triggered as follows: Cln3-Cdk1 phosphorylates the cell-cycle inhibitor Whi5, leading to the nuclear export of phosphorylated Whi5 (de Bruin et al., 2004; Dirick et al., 1995; Schmoller et al., 2015). During G1, the Cln3 signal is integrated by adding phosphates on Whi5, thus the effective time window of Cln3 signals depends on how fast phosphates is to be removed, i.e. the dephosphorylation rate of Whi5 (Liu et al., 2015). Since the dephosphorylated Whi5 will re-enter the nucleus, the faster Whi5 re-enter the nucleus, the shorter the memory will be (Figure 3A).

**Figure 3.**
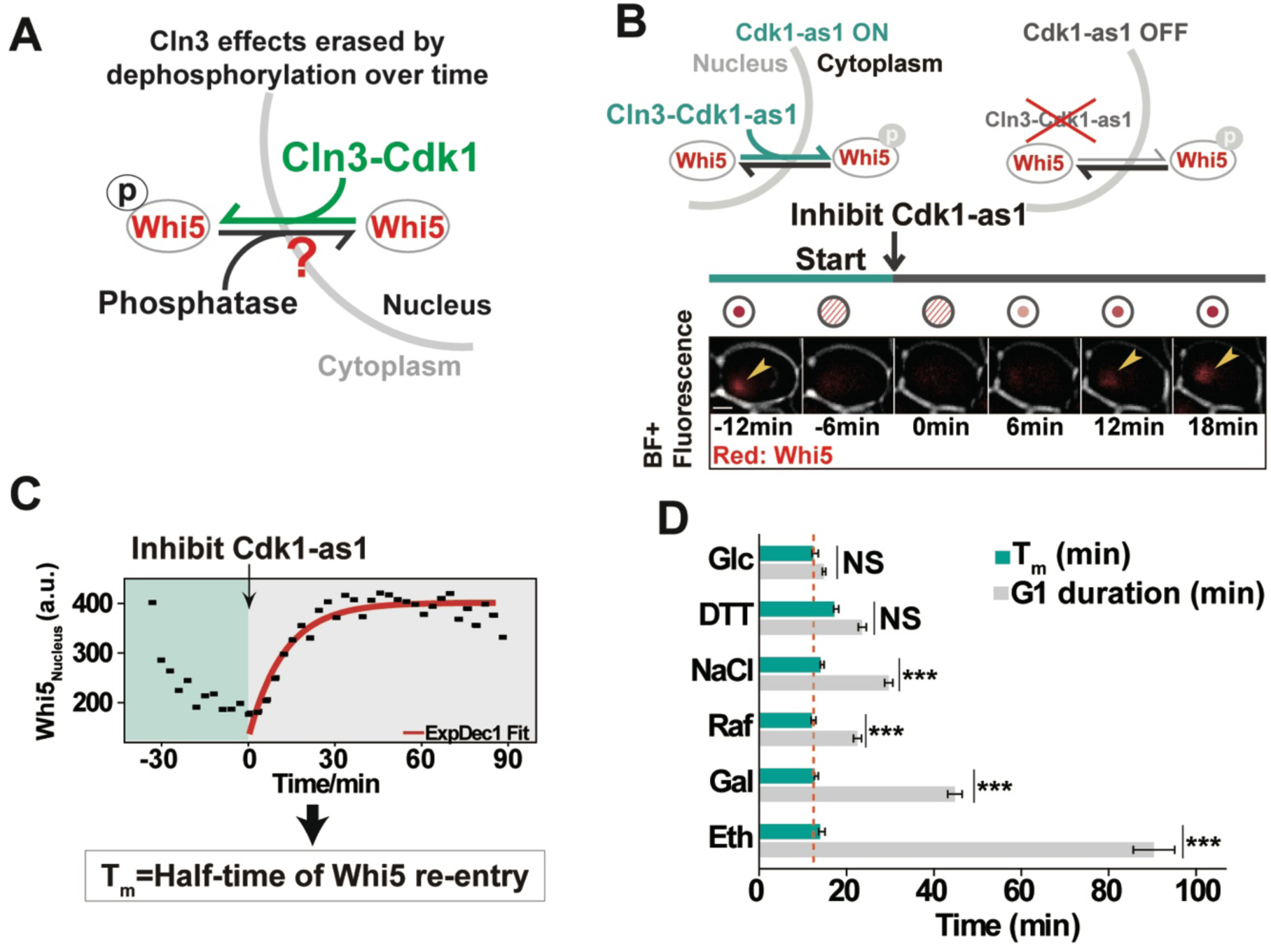
The half-time of nuclear Whi5 re-entery is ∼12 min in a variety of nutrients and stress conditions. (A) Schematic of the regulation of the Whi5 nucleus-cytoplasm translocalization in G1. It is balanced by the phosphorylation and dephosphorylation of Whi5. (B) Schematic of estimating the half-time of nuclear Whi5 re-entry (T_m_) using Cdk1-as1 (top panel). The representative microscopy images of Whi5 dynamics at the indicated time point in response to the Cdk1-as1 activity inhibition right after Whi5 nuclear export (bottom panel). Yellow arrows indicated Whi5. (C) One single cell representative of the Whi5 dynamics in response to the Cdk1-as1 activity inhibition. Tm was estimated using an exponential fit. See Methods. Red curve, exponetial fitting. (D) T_m_ was constant across the different conditions and shorter than the G1 duration under unfavorable conditions. NS, not significant. ***, p value < 0.05.

To measure how long Whi5 re-enters the nucleus, we employed the *Cdc28-as1* allele, whose CDK activity can be inhibited by the ATP analog 1-NM-PP1 (see Methods) (Ubersax et al., 2003). We shut down all CDK activity right after the cell passed the Start and measured how long Whi5 re-entered the nucleus (Figure 3B). We firstly measured the cells in 2% glucose, and the average half-time of Whi5 re-entry is about 12 min (Figure 3C and 3D and Figure S3), which is consistent with our previous results and comparable to the entire G1 length (Liu et al., 2015).

However, as we measured the cells under a variety of nutrient and stress conditions, we found that the half-times of Whi5 re-entry were almost the same, including the unfavorable conditions where cells have a much longer G1 duration (Figure 3D and Figure S3). These results suggested, rather than the entire G1, cells have a relative constant and short time window to integrate environmental cues through Cln3 under various conditions.

## Discussion

Here we propose that effects of Cln3 on triggering Start are not unremittingly accumulated through G1; rather there is a temporary “memory” for Cln3 whereby the cell registers and integrates recent environmental signals to decide whether or not to divide under various conditions.

An intriguing observation of ours is that the short “memory” is more or less independent of nutrient conditions, considering that this memory is for better evaluating the nutrient condition. One possible explanation is that this time scale reflects some kind of relevant time scale in environmental fluctuations in the typical environments yeast cell live. It would be of interest to investigate the memory time of different yeast species living in different natural environments.

Protein phosphorylation is a dynamic and reversible post-translational modification widely implemented in all kinds of biological processes to regulate a myriad of activities including protein activation, translocation, degradation, binding/unbinding and signaling. The memory/integration mechanism studied here could be used in other places. For instance, a similar short-term integration window based on phosphorylation regulation has been discovered during *Drosophila* development (Di Talia and Wieschaus, 2012). Note that the memory time can be easily fine-tuned by the number of phosphorylation sites, binding affinity of phosphatase, or the expression level of phosphatase.

Cells possess memories of different time scales via different mechanisms to solve different problems. Even for the same problem, it may involve different time scales. An example is for cell such as yeast to evaluate the environmental condition, whose fluctuation may involve many time scales, to decide when to divide. Our previous and this study show that yeast cells implement both long (in the order of doubling times 50-220 minutes) and short (∼12 minutes) memories to optimize its decision making (Liu et al., 2015; Qu et al., 2019). We anticipate that many additional cases of short-term and long-term cellular memories exist in order to enable cells to record information about their environment, and understanding the diverse molecular means through which these memories are recorded will be of interest.

## Material and methods

### Yeast strains and plasmids

All the strains used in this study were constructed based on W303. The yeast strains used in this study were listed in Supplemental material Table S1. All the yeast transformations were performed with standard protocols as below. Cells prepared for transformation were grown in YPAD overnight. The overnight cultures were diluted to A_600nm_ of 0.05 into YPAD till OD∼0.8. Pellet cells by centrifugation at 2500 RCF, 5 min at room temperature. Discard supernatant. Wash cells twice in 1/10 volume of 100 mM LiAc; Re-suspend cells in 1/100 volume of 100 mM LiAc (1 transformant / 3 ml culture). Prepare transformation mixture. Distribute 153 μl transformation mixture into each tube with 30 μl cell suspension. Add 1-3 μl plasmid or 20 μl digested plasmid or 20 μl PCR products and mix by vortex briefly. 30 °C incubate 30 min, then 42 °C for 15 min. Centrifuge 1 min at 2500 RCF, re-suspend cell into 100 μl YPAD medium. Plate on selective plates, colonies will show after 2 days, if tag gene or deletion gene or using NAT, G418 selective plates, 30 °C incubate 2-3 hr, then plate on selective plates. Transformation mixture is prepared as below: 15 μl 1 M LiAc, 20 μl (2 mg/ml) carrier DNA, and 18 μl DMSO. Resulting constructs were confirmed by PCR and microscopy.

Plasmids were replicated in DH5α *Escherichia coli*. Plasmids used in this study were listed in Supplemental material Table S2. Standard protocols were used for molecular cloning. Resulting constructs were confirmed by both colony PCR and sequencing.

### Construction of Cln3-controlled system using a light-inducible module

The Cln3-controlled system was constructed using the PhyB-PIF optogenetic module. PhyB is a fragment of *Arabidopsis thaliana* phytochrome B, and PIF is a fragment of phytochrome interaction factor 6. After covalently bound with PCB (a membrane-permeable small-molecule chromophore phycocyanobilin), PhyB and PIF associates with each other very rapidly in response to red (650 nm) light. They dissociate very rapidly in response to infrared (750 nm) light (Levskaya et al., 2009; Shimizu-Sato et al., 2002). We tagged the PhyB with a CAAX (plasma membrane targeting sequence) and tagged Cln3 with 2 yEGFPs and a PIF to control the localization of Cln3 by light. When exposed to red light, PIF-yEGFP-yEGFP-Cln3 fusion protein can be anchored on the plasma membrane. When exposed to infrared light, PIF-yEGFP-yEGFP-Cln3 can be anchored inside the nucleus by the NLS of Cln3 (Miller and Cross, 2000). Therefore, we can not only control the localization of Cln3 but also measure its nuclear concentration in real-time.

The expression of PIF-yEGFP-yEGFP-Cln3 fusion protein is driven by an IPTG-inducible promoter *GlacSpr* (Liu et al., 2015). The Cln3-controlled strain was cultured in the medium containing 0.5 μM IPTG to induce constitutive expression of the light-controlled Cln3. Therefore, the nuclear Cln3 pulse can be generated only by the control of light.

Light power used to generate sharp pulse is 0 w for 650 nm and 4.5 w for 750 nm; light power used to generate longer and lower pulse is 0.5 w for 650 nm and 4.5 w for 750 nm (Figure 2 and S2).

### Growth conditions

For imaging, single colonies were picked from YPAD agar plates and dispensed into 3∼4 ml relative media. Cells were then grown at 30°C overnight in a shaking incubator. The overnight cultures were diluted to OD_600_ of 0.05 into 4ml relative media. Cells were grown to OD_600_ of 0.5 for imaging. The Cln3-controlled strain was cultured with 27 μM PCB for 1-2 hr in dark before imaging to allow the incorporation of the chromophore.

### Microfluidics chips

The microfluidics chips were constructed with polydimethylsiloxane using standard techniques of soft lithography and replica molding. The cells were quickly concentrated, loaded into the microfluidic chip. Syringe filled with 1ml medium was connected to the inlet using soft polyethylene tubing. The flow of medium in the chip was maintained by an auto-controlled syringe pump (TS-1B, Longer Pump Corp., Baoding, China) with a constant velocity of 66.7 μl/hour.

### Time-lapse microscopy and image analysis

The Cln3-controlled cells were taken images by confocal microscope. 256×256 pixel-images were taken by UltraVIEW VoX Laser Confocal Imaging System (PerkinElmer, Watham, MA) and a CSU-X1 spinning disk confocal (Yokogawa, Tokyo, Japan) on a Nikon Ti-E inverted microscope equipped with the APO TIRF 100 × OIL NA 1.45 objective lens, the motorized XY stage and the Perfect-Focus System (Nikon Co., Tokyo, Japan). Images were acquired every 3 min by a Hamamatsu C9100-13 EMCCD (Hamamasu Photonics K. K., Hamamatsu City, Japan) camera. We used 20% 488 nm laser and 20% 561 nm laser for detection of yEGFP and tdTomato proteins, respectively. Exposure time for yEGFP and tdTomato is 150 ms and 100 ms, respectively. *Cdk1-as1* mutant measurements were taken by a Nikon Observer microscope with an automated stage and Perfect-Focus-System (Nikon Co., Tokyo, Japan) using an Apo 100 ×/1.49 oil TIRF objective. Cell segmentation and fluorescent quantification were performed by Cellseg as previously described (Liu et al., 2015). The Cln3*_Nuclear_ intensity was estimated as the mean intensity of the 5×5 brightest yEGFP-Cln3 pixels in the cell and the Whi5_Nuclear_ intensity was estimated as the mean intensity of the 5×5 brightest Whi5-tdTomato pixels in the cell.

### Estimate the half-time of Whi5 re-activation

We used the inhibitor-sensitive *Cdk1-as1* mutant (Ubersax et al., 2003) to estimate how long effects of Cln3 on Whi5 inactivation can last. The kinase activity of *Cdk1-as1* was inhibited by 1-NM-PP1 (EMD Millipore, CAS# 529581) with the concentration of 25 μM.

## Acknowledgments

The authors thank Kyle M. Loh for insightful discussions and comments on the manuscript. This work was partly supported by The National Natural Science Foundation of China (NSFC31700733) and The National Key Research and Development Program of China (2018YFA0900700).

## Author contributions

C.T. X.Y. and Y.Q. designed the project. Y.Q. and J.J. designed, performed the experiments, and analyzed the data. X.L. wrote computer programs and analyzed the data. C.T. and X.Y. supervised the whole project; Y.Q., J.J., C.T. and X.Y. wrote the paper.

## Supplementary figures

**Figure S1.**
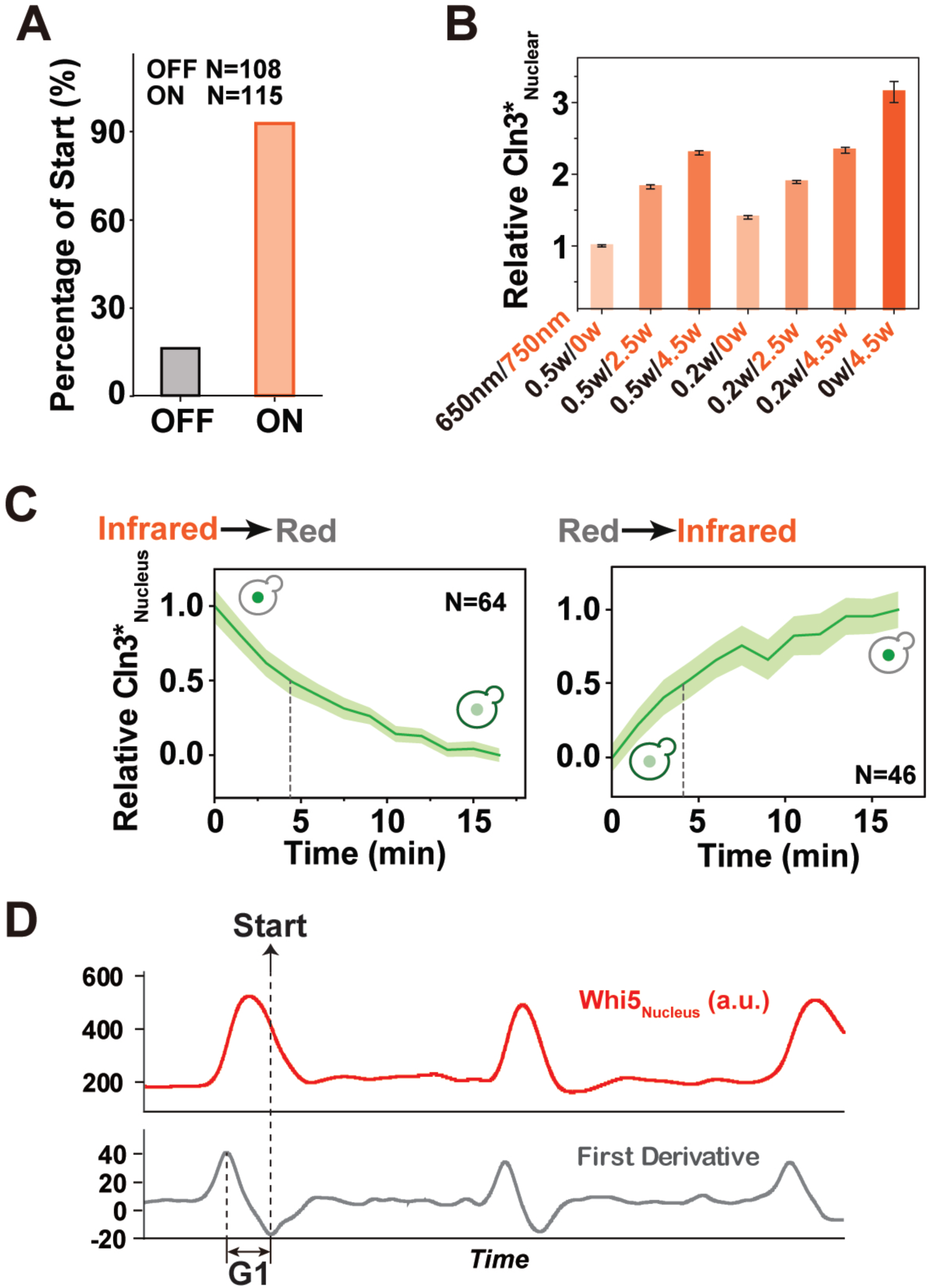
Construction of the Cln3-controlled system. (A) Percentage of Start in cells exposed to red light (OFF state) and infrared light (ON state) during 2.5 h observation. Cells were arrested in G1 when exposed to red light (OFF state) and went through cell cycle normally when exposed to infrared light (ON state). (B) Averaged relative nuclear Cln3* intensity in cells exposed to the indicated light intensity. (C) Real-time observation of dynamics of relative nuclear Cln3* intensity in cells in response to the red light (left panel) and infrared light (right panel). Data were shown as mean ± SEM. (D) Defination of the Start point by the nuclear Whi5 dynamics. The lowest first derivative of the nuclear Whi5 curve was defined as the Start point where yeast committed to the cell cycle. The G1 duration was defined as the time during which Whi5 stays in the nucleus (time between the highest first derivative and the lowest first derivation).

**Figure S2.**
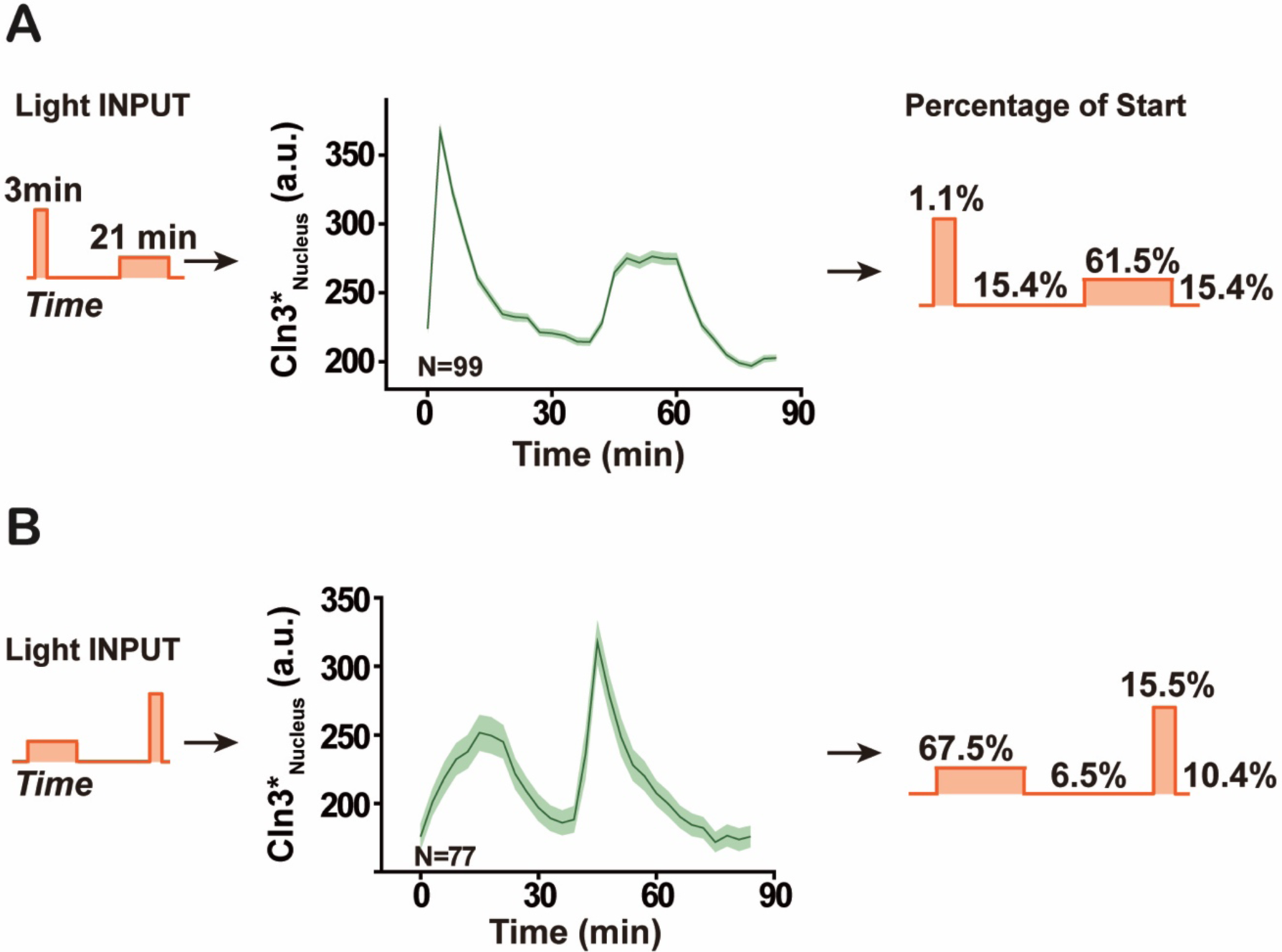
Start depends on the time duration of the nuclear Cln3* signal instead of the instantaneous nuclear Cln3* intensity. (A-B) Real-time observation of dynamics of nuclear Cln3* intensity in cells (middle panel) exposed to the indicated light input (left panel). Data were shown as mean ± SEM. The percentage of cells that passed through Start induced by the indicated Cln3* pulse (right panel). See Methods for light power.

**Figure S3.**
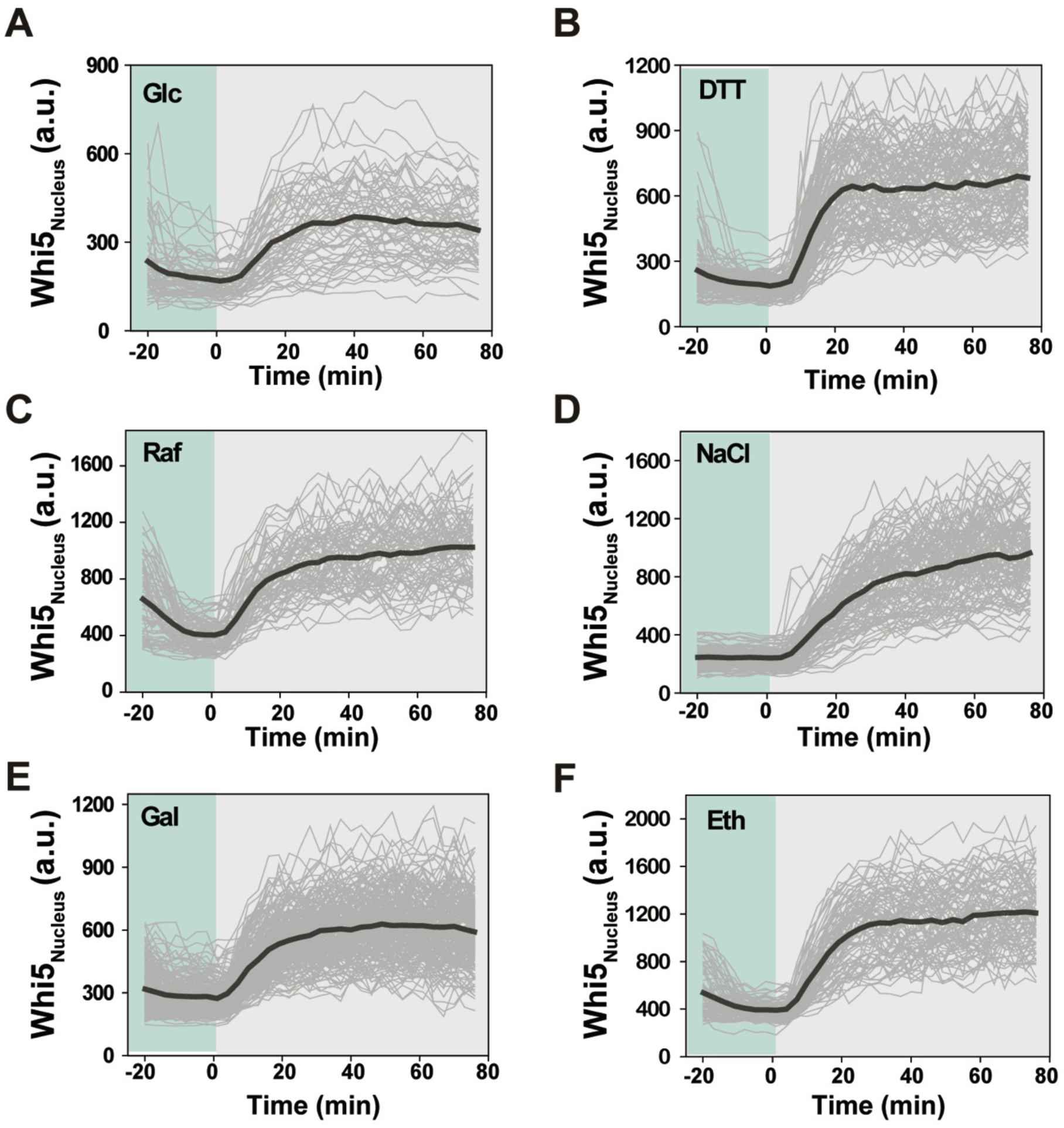
Real-time observation of dynamics of nuclear Whi5 intensity in cells under different conditions (A-F). The black curve indicated the mean. Each light gray curve indicated a single cell data. Green shading represented the time before the inhibition of Cdk1-as1. Gray shading represented the time after the inhibition of Cdk1-as1.

## References

Berg, H.C. (2000). Motile behavior of bacteria. Phys Today 53, 24–29.

Burrill, D.R., and Silver, P.A. (2010). Making cellular memories. Cell 140, 13–18.

Cai, Y., and Futcher, B. (2013). Effects of the yeast RNA-binding protein Whi3 on the half-life and abundance of CLN3 mRNA and other targets. PLoS One 8, e84630.

Cross, F.R., and Blake, C.M. (1993). The yeast Cln3 protein is an unstable activator of Cdc28. Mol Cell Biol 13, 3266–3271.

Csermely, P., Kunsic, N., Mendik, P., Kerestely, M., Farago, T., Veres, D.V., and Tompa, P. (2020). Learning of Signaling Networks: Molecular Mechanisms. Trends Biochem Sci 45, 284–294.

de Bruin, R.A., McDonald, W.H., Kalashnikova, T.I., Yates, J., 3rd, and Wittenberg, C. (2004). Cln3 activates G1-specific transcription via phosphorylation of the SBF bound repressor Whi5. Cell 117, 887–898.

Di Talia, S., and Wieschaus, E.F. (2012). Short-term integration of Cdc25 dynamics controls mitotic entry during Drosophila gastrulation. Dev Cell 22, 763–774.

Dirick, L., Bohm, T., and Nasmyth, K. (1995). Roles and regulation of Cln-Cdc28 kinases at the start of the cell cycle of Saccharomyces cerevisiae. EMBO J 14, 4803–4813.

Doncic, A., Falleur-Fettig, M., and Skotheim, J.M. (2011). Distinct interactions select and maintain a specific cell fate. Mol Cell 43, 528–539.

Epstein, C.B., and Cross, F.R. (1994). Genes that can bypass the CLN requirement for Saccharomyces cerevisiae cell cycle START. Mol Cell Biol 14, 2041–2047.

Gallego, C., Gari, E., Colomina, N., Herrero, E., and Aldea, M. (1997). The Cln3 cyclin is down-regulated by translational repression and degradation during the G1 arrest caused by nitrogen deprivation in budding yeast. EMBO J 16, 7196–7206.

Gao, Z., Sun, H., Qin, S., Yang, X., and Tang, C. (2018). A systematic study of the determinants of protein abundance memory in cell lineage. Science Bulletin 63, 1051–1058.

Gardner, T.S., Cantor, C.R., and Collins, J.J. (2000). Construction of a genetic toggle switch in Escherichia coli. Nature 403, 339–342.

Gaydos, L.J., Wang, W., and Strome, S. (2014). Gene repression. H3K27me and PRC2 transmit a memory of repression across generations and during development. Science 345, 1515–1518.

Hall, D.D., Markwardt, D.D., Parviz, F., and Heideman, W. (1998). Regulation of the Cln3-Cdc28 kinase by cAMP in Saccharomyces cerevisiae. EMBO J 17, 4370–4378.

Harvey, Z.H., Chen, Y., and Jarosz, D.F. (2018). Protein-Based Inheritance: Epigenetics beyond the Chromosome. Mol Cell 69, 195–202.

Johnson, A., and Skotheim, J.M. (2013). Start and the restriction point. Curr Opin Cell Biol 25, 717–723.

Jorgensen, P., and Tyers, M. (2004). How cells coordinate growth and division. Curr Biol 14, R1014–1027.

Levskaya, A., Weiner, O.D., Lim, W.A., and Voigt, C.A. (2009). Spatiotemporal control of cell signalling using a light-switchable protein interaction. Nature 461, 997–1001.

Liu, X., Wang, X., Yang, X., Liu, S., Jiang, L., Qu, Y., Hu, L., Ouyang, Q., and Tang, C. (2015). Reliable cell cycle commitment in budding yeast is ensured by signal integration. Elife 4.

Ma, W., Trusina, A., El-Samad, H., Lim, W.A., and Tang, C. (2009). Defining network topologies that can achieve biochemical adaptation. Cell 138, 760–773.

Miller, M.E., and Cross, F.R. (2000). Distinct subcellular localization patterns contribute to functional specificity of the Cln2 and Cln3 cyclins of Saccharomyces cerevisiae. Mol Cell Biol 20, 542–555.

Monod, J., and Jacob, F. (1961). Teleonomic mechanisms in cellular metabolism, growth, and differentiation. Cold Spring Harb Symp Quant Biol 26, 389–401.

Novick, A., and Weiner, M. (1957). Enzyme Induction as an All-or-None Phenomenon. Proc Natl Acad Sci U S A 43, 553–566.

Papagiannakis, A., Niebel, B., Wit, E.C., and Heinemann, M. (2017). Autonomous Metabolic Oscillations Robustly Gate the Early and Late Cell Cycle. Mol Cell 65, 285–295.

Parviz, F., Hall, D.D., Markwardt, D.D., and Heideman, W. (1998). Transcriptional regulation of CLN3 expression by glucose in Saccharomyces cerevisiae. J Bacteriol 180, 4508–4515.

Perez, M.F., and Lehner, B. (2019). Intergenerational and transgenerational epigenetic inheritance in animals. Nat Cell Biol 21, 143–151.

Polymenis, M., and Schmidt, E.V. (1997). Coupling of cell division to cell growth by translational control of the G1 cyclin CLN3 in yeast. Genes Dev 11, 2522–2531.

Qu, Y., Jiang, J., Liu, X., Wei, P., Yang, X., and Tang, C. (2019). Cell Cycle Inhibitor Whi5 Records Environmental Information to Coordinate Growth and Division in Yeast. Cell Rep 29, 987–994 e985.

Schaefer, S., and Nadeau, J.H. (2015). The Genetics of Epigenetic Inheritance: Modes, Molecules, and Mechanisms. Q Rev Biol 90, 381–415.

Schmoller, K.M., Turner, J.J., Koivomagi, M., and Skotheim, J.M. (2015). Dilution of the cell cycle inhibitor Whi5 controls budding-yeast cell size. Nature 526, 268–272.

Schneider, B.L., Zhang, J., Markwardt, J., Tokiwa, G., Volpe, T., Honey, S., and Futcher, B. (2004). Growth rate and cell size modulate the synthesis of, and requirement for, G1-phase cyclins at start. Mol Cell Biol 24, 10802–10813.

Shimizu-Sato, S., Huq, E., Tepperman, J.M., and Quail, P.H. (2002). A light-switchable gene promoter system. Nat Biotechnol 20, 1041–1044.

Tu, Y. (2013). Quantitative modeling of bacterial chemotaxis: signal amplification and accurate adaptation. Annu Rev Biophys 42, 337–359.

Tyers, M., Tokiwa, G., Nash, R., and Futcher, B. (1992). The Cln3-Cdc28 kinase complex of S. cerevisiae is regulated by proteolysis and phosphorylation. EMBO J 11, 1773–1784.

Ubersax, J.A., Woodbury, E.L., Quang, P.N., Paraz, M., Blethrow, J.D., Shah, K., Shokat, K.M., and Morgan, D.O. (2003). Targets of the cyclin-dependent kinase Cdk1. Nature 425, 859–864.

Verges, E., Colomina, N., Gari, E., Gallego, C., and Aldea, M. (2007). Cyclin Cln3 is retained at the ER and released by the J chaperone Ydj1 in late G1 to trigger cell cycle entry. Mol Cell 26, 649–662.

Wang, H., Carey, L.B., Cai, Y., Wijnen, H., and Futcher, B. (2009). Recruitment of Cln3 cyclin to promoters controls cell cycle entry via histone deacetylase and other targets. PLoS Biol 7, e1000189.

Yaglom, J., Linskens, M.H., Sadis, S., Rubin, D.M., Futcher, B., and Finley, D. (1995). p34Cdc28-mediated control of Cln3 cyclin degradation. Mol Cell Biol 15, 731–741.

Yahya, G., Parisi, E., Flores, A., Gallego, C., and Aldea, M. (2014). A Whi7-anchored loop controls the G1 Cdk-cyclin complex at start. Mol Cell 53, 115–126.

Yang, X., Jost, A.P., Weiner, O.D., and Tang, C. (2013). A light-inducible organelle-targeting system for dynamically activating and inactivating signaling in budding yeast. Mol Biol Cell 24, 2419–2430.

